# Natural variation identifies a *Pxy* gene controlling root vascular organization and formation of nodules and lateral roots in *Lotus japonicus*

**DOI:** 10.1101/2020.04.08.031260

**Authors:** Yasuyuki Kawaharada, Niels Sandal, Vikas Gupta, Haojie Jin, Maya Kawaharada, Korbinian Schneeberger, Jens Stougaard, Stig U. Andersen

## Abstract

Forward and reverse genetics using the model legumes *Lotus japonicus* and *Medicago truncatula* have been instrumental for identifying the essential genes governing legume-rhizobial symbiosis. However, little is known about the effects of intraspecific variation on symbiotic signaling. The *Lotus* accessions Gifu and MG20 show differentiated phenotypic responses to the *Mesorhizobium loti exoU* mutant that produces truncated exopolysaccharides. Using Quantitative Trait Locus sequencing (QTL-seq), we identify the *Pxy* gene as a component of this differential *exoU* response. *Lotus Pxy* encodes a leucine-rich-repeat kinase similar to *Arabidopsis* PXY, which regulates stem vascular development. We show that *Lotus pxy* insertion mutants display defects in root vascular organization, as well as lateral root and nodule formation. Our work links *Pxy* to *de novo* organogenesis in the root, highlights the genetic overlap between regulation of lateral root and nodule formation, and demonstrates that specific natural variants of *Pxy* differentially affect nodulation signaling.

## Introduction

The symbiotic interaction between legumes and rhizobia results in the development of new organs called root nodules. These symbiotic organs are established by two tightly synchronized processes, bacterial infection and nodule organogenesis. Rhizobia sense specific flavonoids from host legume plants, and in response rhizobia synthesize and secrete lipochitooligosaccharides (Nod factors) that serve as symbiotic signal molecules^1^. In the model legume plant *Lotus japonicus* (*Lotus*), NFR1, NFR5 and NFRe LysM receptors perceive rhizobial Nod factors^2–4^. In subsequent steps NFR5 interacts with the SYMRK LRR receptor and a cytoplasmic kinase NiCK4 that accelerates downstream signaling^5,6^. Further downstream in the symbiosis pathway operating in *Lotus* and/or *Medicago truncatula* (*Medicago*), nucleoporins (NUP85, NUP133, and NENA), cation channels (CASTOR and POLLUX) and calcium channels (CNGC) are involved in releasing the perinuclear calcium spiking^7–14^. This calcium fluctuation is recorded by calcium calmodulin-dependent protein kinases (CCaMK)^15–18^, and activated CCaMK phosphorylates the CYCLOPS transcription factor^19–22^. Additional transcription factors (NSP1, NSP2, ERN1 and NIN) are then required for infection thread formation^23–29^ involving a pectate lyase encoded by *Npl*, actin rearrangement by NAP1, PIR1, SCARN, ARPC1 and DREPP, an E3 ubiquitin ligase encoded by *Cerberus*, an atypical receptor kinase encoded by *RinRK*, a subunit of mediator complex encoded by *Lan,* and novel functions encoded by *Rpg, Vapyrin* and *CBS*^30–40^. The simultaneous development of nodule organs is mediated by NSP1, NSP2, ERN1, NIN and NF-Ys transcription factors^41,42^ that together with localized changes in plant hormone homeostasis (cytokinin, auxin and jasmonic acid) regulate initiation of infection thread formation and cell divisions leading to nodule organogenesis^43–46^. In contrast, ethylene and abscisic acid suppress nodule formation^47–49^.

Distinct mutant phenotypes and monogenic inheritance is the foundation for the characterization of this central pathway that is highly conserved across model legumes and other symbiotic interactions, for example the Frankia-actinorhizal symbiosis, highlighting the importance of the core components of the symbiosis pathway^50^. Inter- and intraspecific differences reflecting natural variation have, however, also been found. One such example is the influence of rhizobial exopolysaccharides on root nodule development, primarily infection thread formation. Classical genetic and biochemical studies using the *Lotus* Gifu accession have shown that the EPR3 receptor perceives and recognizes EPS produced by *M. loti* and controls infection thread formation^51,52^. Interestingly, variability in exopolysaccharide responses towards truncated exopolysaccharide synthesized by the *M. loti exoU* mutant was found among 65 *Lotus* natural accessions^53^. Like Gifu, 15 wild *Lotus* accessions formed only white, uninfected nodule primordia. A less stringent response was observed with 45 accessions where up to 50 % of the plants developed at least one pink infected nodule, and an intermediate response was found in eight accessions, including MG20, that developed pink nodules on 50 -75 % of the plants. Such quantitative phenotypic differences suggest that there are complex multigene traits involved in diversification that can be uncovered using genome-wide genetic methodologies.

Here, we use Quantitative Trait Locus sequencing (QTL-seq) to identify *Pxy* as a causal component of the differential *exoU* response and show that *Pxy* is required for normal root vascular organization and lateral root and nodule formation in *Lotus*.

## Results

### *Lotus japonicus* ecotypes Gifu and MG20 show contrasting symbiotic phenotypes on inoculation with the *Mesorhizobium loti exoU* mutant

Previous studies described a differential, quantitative symbiotic response to the *M. loti exoU* mutant among *Lotus* accessions, suggesting that the *exoU* response could be a multigene trait^53^. To investigate this further, we focused on two accessions, Gifu and MG20^54,55^, which showed clearly contrasting phenotypes. When inoculated with the *exoU* mutant Gifu showed impaired infection thread formation and formed small uninfected nodules (nodule primordia), while MG20 developed pink effective nodules. Approximately 78% of the MG20 plants formed 1 to 6 pink, nitrogen-fixing nodules per plant at 6 weeks post inoculation, while Gifu plants did not form any pink nodules (**Fig. 1**).

**Figure 1.**
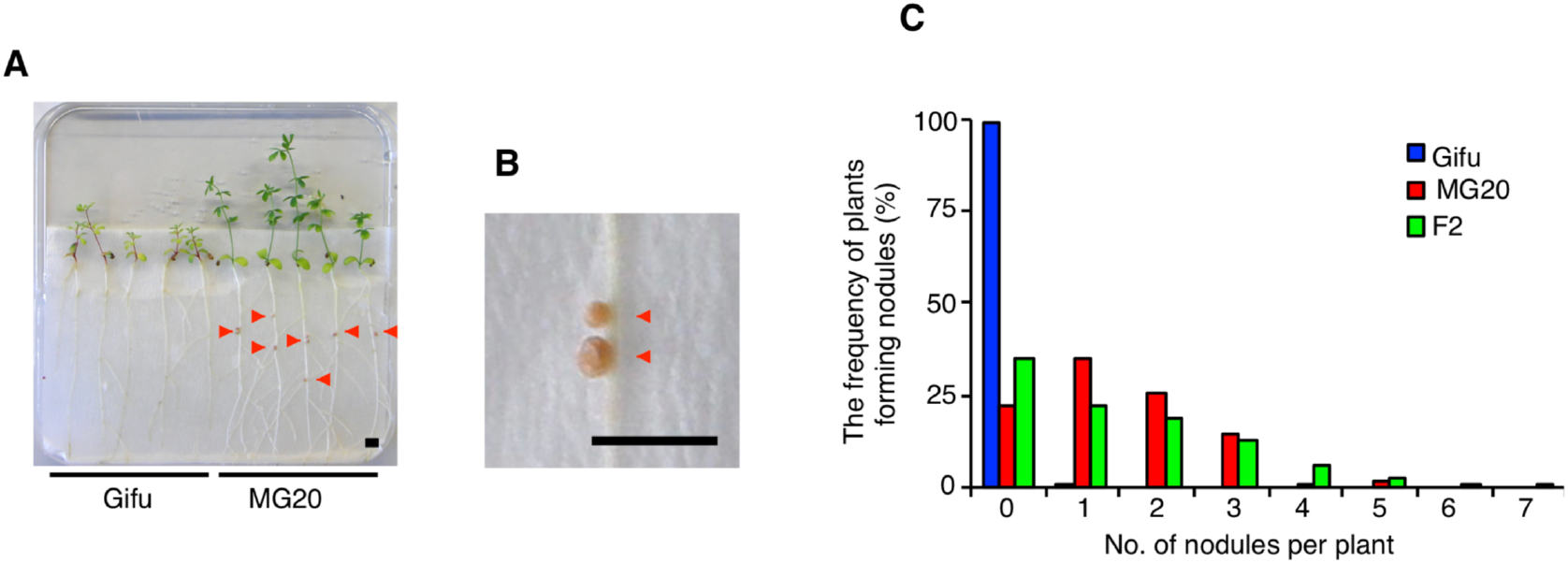
*M. loti exoU* response in *Lotus* ecotypes Gifu, MG20 and F2 progeny. **A**: The symbiotic phenotypes of Gifu and MG20 plants 6 weeks post-inoculation with the *M. loti exoU* mutant. **B**: Infected pink nodules in MG20 inoculated with the *exoU* mutant. **C**, Frequency distribution of plants forming pink nodules in Gifu, MG20 and F2 progeny derived from the cross between Gifu and MG20 6 weeks post-inoculation with the *exoU* mutant. *n* = 98 (Gifu), 125 (MG20), and 319 (F2 progeny). Red arrows show pink nodules in MG20 plants. The bar is 0.5 cm.

As a first attempt to investigate the inheritance of the *exoU* response, we examined a Gifu x MG20 F2 population. Approximately 65% of the F2 plants formed nodules with an intermediate frequency, indicating that the trait is not monogenic (**Fig. 1C**). Genotyping the F2 plants that formed at least three pink nodules using SSR markers^56,57^, we identified two chromosomal regions associated with pink nodule formation, one at the end of chromosome 3 and one at the middle of chromosome 4 **(Fig. 2A**). These results suggested that the difference in *exoU* nodulation phenotype between Gifu and MG20 is genetically controlled and that the number of pink nodules is likely determined by multiple genes.

**Figure 2.**
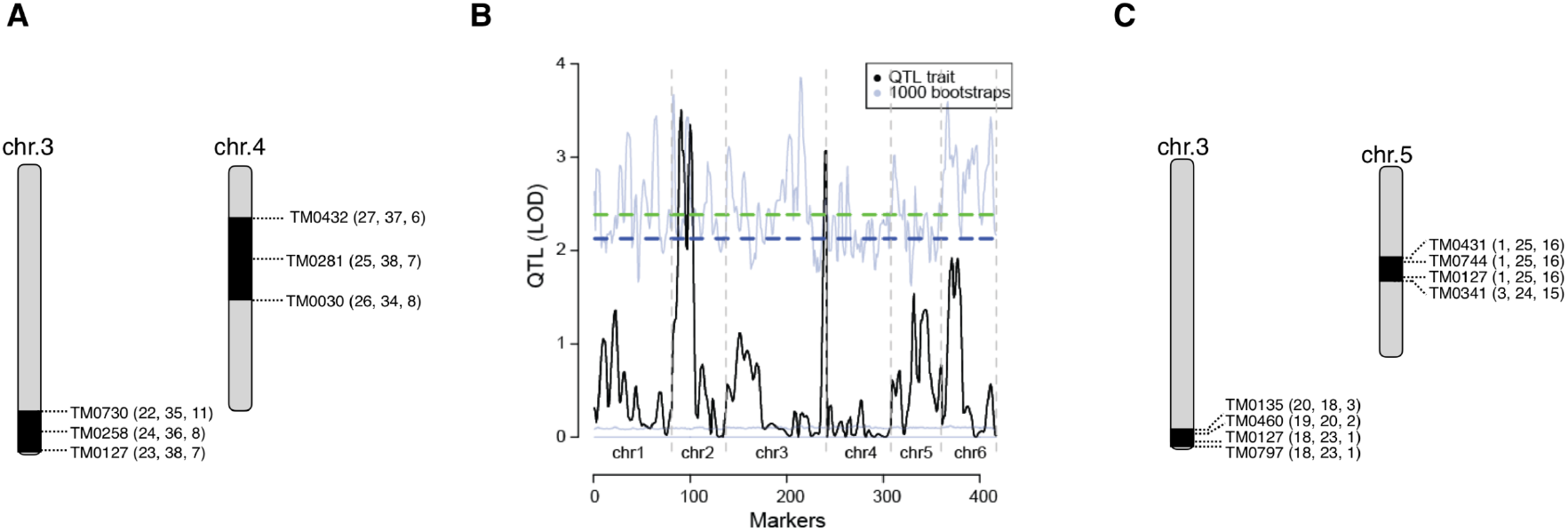
QTL-seq mapping of the *exoU* response. **A**: Linkage analysis of symbiotic phenotypes in F2 plants derived from the cross between Gifu and MG20. Plants forming more than 3 pink nodules after inoculation with the *exoU* mutant were selected and genotyped. **B**: QTL analysis of *exoU* response based on analysis of Gifu x MG20 RILs. The quantitative trait scored was the average number of nodules after *exoU* inoculation. 5 and 10% false discovery rate levels are indicated by green and blue dashed lines, respectively. Light blue lines delimit the results of 1000 QTL analyses on permuted data, which were used to determine the significance thresholds. **C**: A linkage analysis of symbiotic phenotypes in F2 plants derived from the cross between Gifu and RI34, which is one of the RILs forming pink nodules after *exoU* inoculation. Black areas identify candidate regions. **A, C:** Numbers next to the SSR markers indicate numbers of individuals with MG20, heterozygous and Gifu genotypes, respectively.

### QTL analysis using recombinant inbred lines

In order to substantiate the F2 segregation analysis, we characterized the symbiotic phenotype of 114 recombinant inbred lines (RILs) derived from a Gifu x MG20 cross^56,58^, counting the number of *exoU*-induced nodules per plant (**Supplemental table 1**). The symbiotic phenotype and nodulation score were then analyzed using R/qtl^59^, and we detected three chromosomal regions associated with the number of pink nodules on chromosomes 2 and 3 (**Fig. 2B**). To further narrow down these regions we used RI-34, which has an MG20 genotype at the top of chromosome 2 and at the end of chromosome 3 and formed pink nodules with the *exoU* mutant (**Supplemental figure 1**). F2 plants of RI-34 back-crossed to Gifu were tested for symbiotic phenotype and genotyped. These F2 plants showed segregation for pink nodule formation, and a linkage-analysis using SSR markers was therefore performed. Using plants forming at least three pink nodules, two genomic regions were detected (**Fig. 2C**). The region at the end of chromosome 3 was enriched for MG20 alleles, while the region at the center of chromosome 5 showed enrichment of Gifu alleles (**Fig. 2C**).

### QTL-seq fine-mapping of candidate genes on chromosome three

In order to gain further genetic resolution, we took a QTL-seq approach and sequenced bulked segregant pools. To establish the bulks, a large MG20 x Gifu F2 population inoculated with the *exoU* mutant was scored for the number of pink nodules and divided into two pools with contrasting phenotypes for collection of genomic DNA. Pool 1 consisted of 450 plants that formed at least three pink nodules and pool 2 comprised 6,801 plants that did not develop any pink nodules. Another 2,515 plants with intermediate phenotypes were discarded (**Table 1**). Genomic DNA was extracted from pools 1 and 2 and sequenced on an Illumina Hi-Seq 2000 instrument, resulting in more than 100 million reads and 20 x genome coverage per pool (**Supplemental table 2**).

**Table 1.**
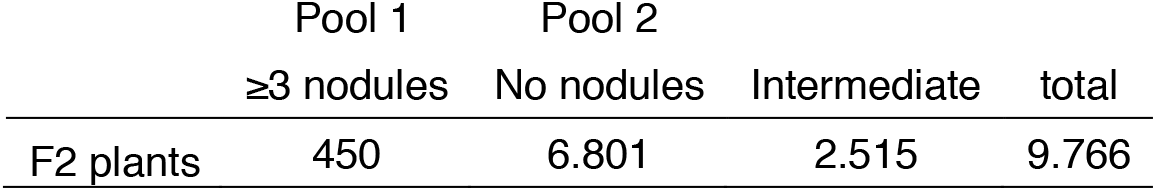
Segregation into extreme phenotypic pools for QTL-seq.

**Table 2.**
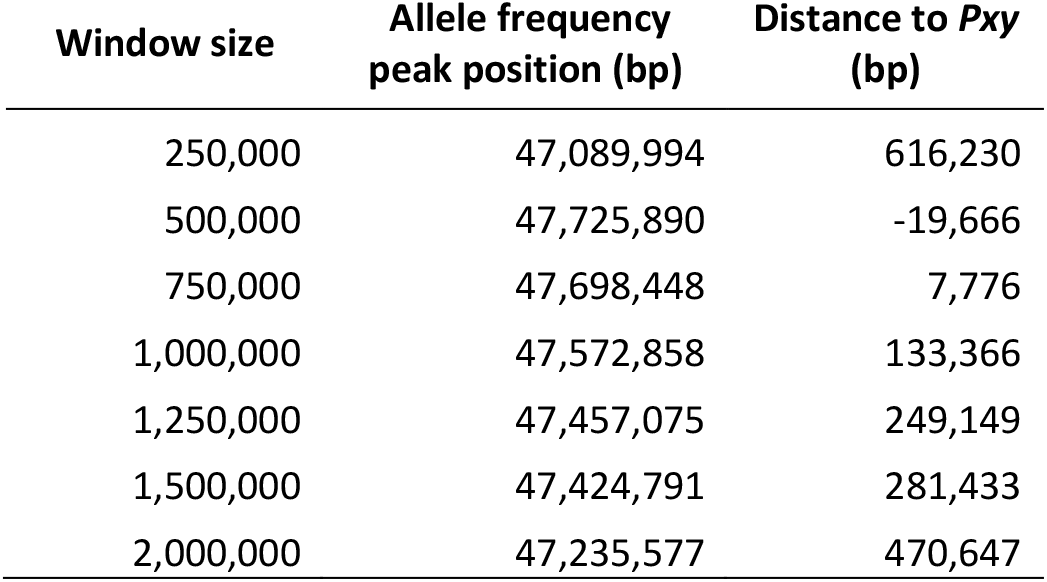
Peak estimates for the exoU nodulation QTL on chromosome 3. See also Figure 3.

We calculated allele frequencies for the two pools at all polymorphic positions and used this as a basis for sliding window analysis of allele frequency differences between the two pools. Focusing on the bottom of chromosome 3, which was also robustly detected by the mapping approaches based on geno- and phenotyping of individual plants (**Fig. 2**), we found a clear increase in allele frequency differences towards the end of the chromosome. The pool with at least three pink *exoU* nodules showed enrichment for MG20 alleles. This signal was detected regardless of the window size used (**Fig. 3**) and it is consistent with the RIL and F2 mapping results (**Fig. 2**).

**Figure 3.**
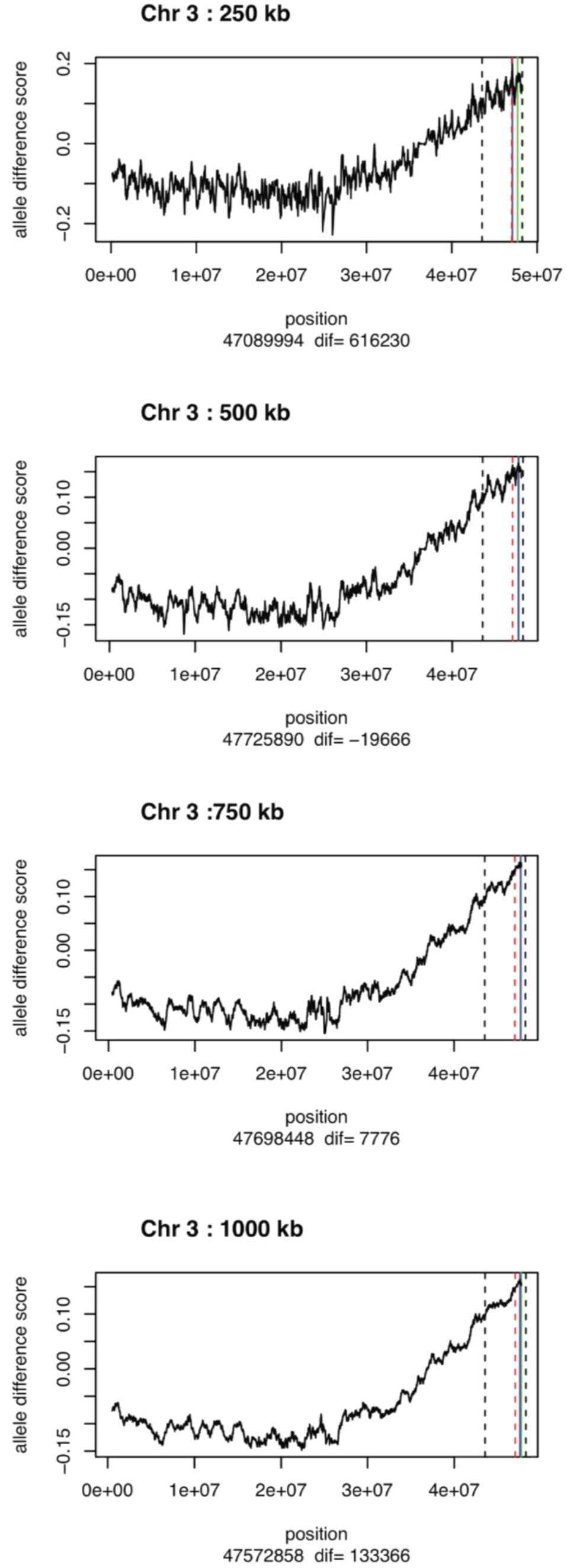
Identification of a QTL peak on chromosome 3 by QTL-seq analysis. Different sliding window sizes were used for QTL peak estimates on chromosome 3. The solid blue line indicates the peak estimate. The solid green line shows the position of the *Pxy* gene, whereas the red dashed lines indicate the genomic position of the closest linked marker according to the RIL-based QTL analysis. The leftmost dashed black lines show the position of the neighboring flanking marker from the QTL analysis. The rightmost black lines indicate the position of the last polymorphic SNP marker in the QTL-seq analysis. The exact position of the estimated peak and the distance (dif) between the peak estimate and the *Pxy* gene (blue and green lines) are indicated below the plot.

We then examined the results of the sliding window analyses in greater detail to accurately estimate the position of the allele frequency difference peak, at which the causal gene(s) should be located^60^. Disregarding the plots with a 250 kb window size, which appeared too noisy, there was a clear trend in the allele frequency difference peak estimates - the larger the window size, the more the peak estimate shifted away from the chromosome end (**Table 2**). The reason for this trend is most likely the close proximity of the QTL to the end of the chromosome, where the last polymorphic SNP marker is found at 48,26 Mb, making it impossible for large sliding windows to detect peaks at the extreme ends of the chromosome. Since the peak estimates of the 500 and 750 kb windows were remarkably similar, we chose their average as the final peak position estimate of 47,712,169 kb (**Table 2**).

### *Pxy* is a causal component of the differential *exoU* response

Sorting the candidates according to their peak distance identified a phloem intercalated with xylem (*Pxy,* chr3.CM0261.600.r2.a in the *Lotus* MG20 genome assembly v2.5) and a Xyloglucan endotransglucosylase/hydrolase 2 (*Xdh2,* chr3.CM0261.560.r2.a) as the closest nearby genes harboring Gifu/MG20 polymorphisms (**Supplemental table 3**). Upon closer manual inspection of the region, an unannotated gene was also found. It was similar to *Structural maintenance of chromosomes 6* (*Smc6*) from *Arabidopsis* and harbored a large number of polymorphisms (**Supplemental table 3**). To carry out complementation tests, these three genes or their coding sequences from Gifu and MG20 were fused to the *Lotus* ubiquitin promoter and transformed into Gifu or MG20 plants using *Agrobacterium rhizogenes*^61–63^. The resulting transgenic hairy roots were inoculated with the *exoU* mutant and the symbiotic phenotype was scored. Regardless of the construct used, the transformed Gifu roots did not consistently form nodules, whereas the MG20 roots did (**Table 3 and Supplemental table 4**).

**Table 3.** *Pxy* complementation experiment results.

| Plant | Vector | Total plants | No. of plants forming nodules | Total nodules | Pink nodule | White nodule | % (No. of plants forming nodules/total plants) |
| --- | --- | --- | --- | --- | --- | --- | --- |
| Gifu | pIV10 | 84 | 4 | 4 | 0 | 4 | 4.7 |
|  | pUbi::MG20<br>*Pxy* | 90 | 5 | 6 | 0 | 6 | 5.6 |
|  | pUbi::Gifu<br>*Pxy* | 34 | 2 | 3 | 0 | 3 | 5.9 |
| MG20 | pIV10 | 114 | 23 | 29 | 10 | 19 | 20.2 |
|  | pUbi::MG20<br>*Pxy* | 39 | 12 | 19 | 9 | 10 | 30.8 |
|  | pUbi::Gifu<br>*Pxy* | 39 | 8 | 14 | 8 | 6 | 20.5 |
| RI94 | pIV10 | 103 | 0 | 0 | 0 | 0 | 0 |
|  | pUbi::MG20<br>*Pxy* | 110 | 22 | 26 | 10 | 16 | 20 |
|  | pUbi::Gifu<br>*Pxy* | 55 | 3 | 3 | 1 | 2 | 5.4 |

Considering the multigene inheritance suggested by the segregation analyses, Gifu x MG20 RILs RI-44, 56, 91, 94, 115, 120, 140 and 146 were then selected as recipients for candidate gene transformation (**Table 3 and Supplemental table 4**). These RILs show a Gifu genotype between markers TM0091 and TM0127 in the chromosome 3 candidate region, and could not form any pink nodules when inoculated with *exoU*. Among these RILs, only RI-94 transformed with *Pxy* from MG20 consistently formed nodules when inoculated with *exoU*, while the empty vector and *Xth2* and *Smc6* from MG20 as well as *Pxy* from Gifu transformed into RI-94 resulted in formation of no or very few nodules (**Table 3 and Supplemental table 4**).

The results of these complementation tests indicate that the *Pxy* gene from MG20 is sufficient to allow development of pink nodules with the *exoU* mutant when transferred to a RIL containing the Gifu version of the gene, and that *Pxy* therefore represents a causal component of the differential Gifu/MG20 response to *exoU* inoculation. The *Pxy* gene encodes a leucine-rich repeat receptor kinase protein. The *Lotus* PXY amino acid sequence shares 65 % identity and 78 % similarity with *Arabidopsis thaliana* PXY, and *Lotus* PXY clusters with *Arabidopsis* PXY in a phylogenetic tree of CLE peptide receptors (**Fig. 4A**). In the 1,976 bp putative *Pxy* promoter region, there are twelve SNPs and one base pair deletion between Gifu and MG20. The *Pxy* genes in Gifu and MG20 have three non-synonymous SNPs in exonic regions and three SNPs and two deletions in intronic regions (**Fig. 4B, C**). One of the polymorphisms is found within the LRR5 region of *Lotus* PXY, where MG20 has an arginine in contrast to the Gifu glycine residue, which is the more common variant (**Fig. 4D**).

**Figure 4.**
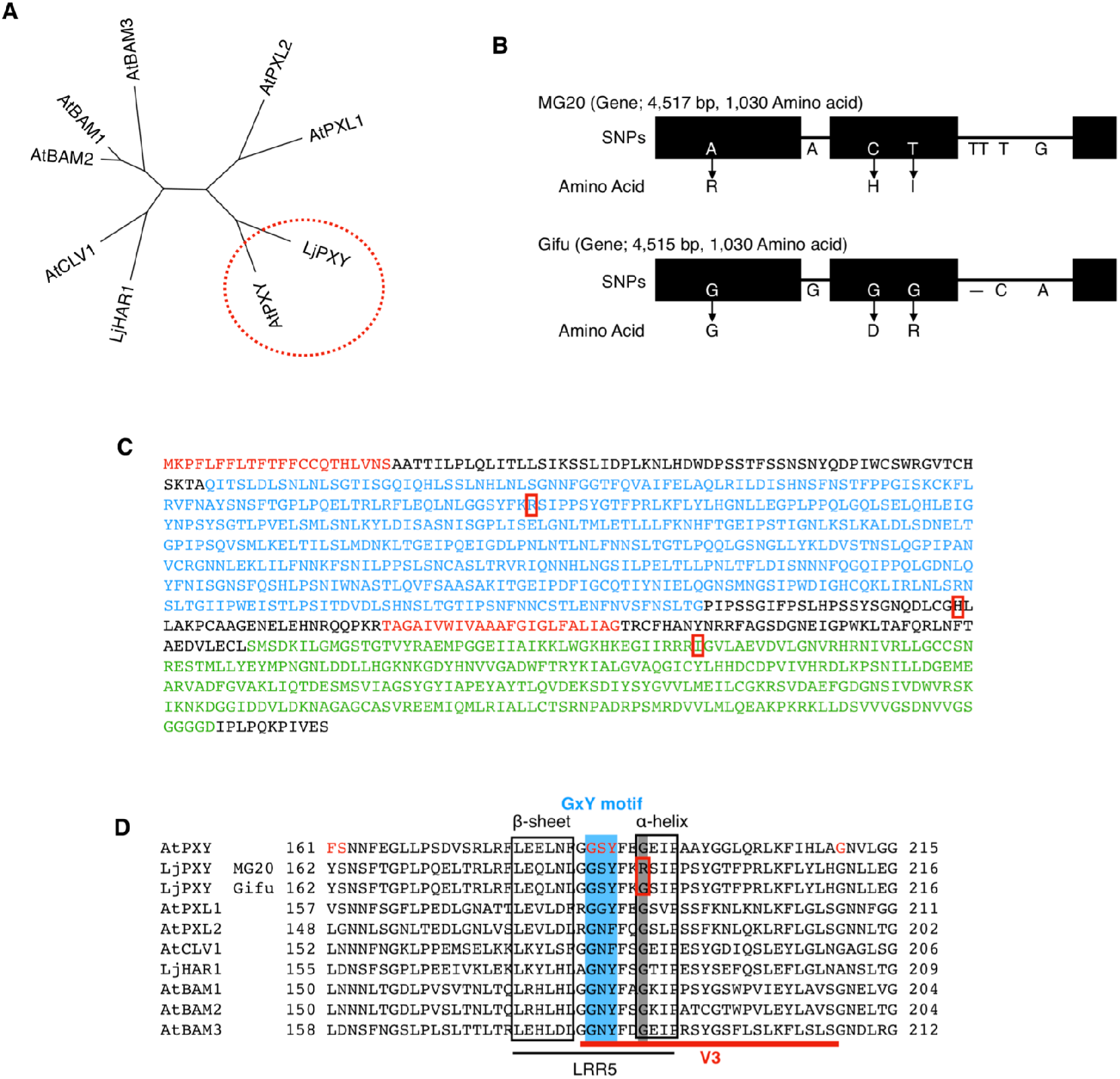
Gifu and MG20 PXY sequences. **A**: Neighbor-joining phylogenetic tree of CLE peptide receptors in *Lotus* and *Arabidopsis*. **B**: Schematic of the *Pxy* gene structure in Gifu and MG20. Eight nucleotides in *Pxy* are different between Gifu and MG20 and three of them cause amino acid changes. **C**: PXY amino acid sequence in MG20. Signal peptide and trans-membrane domains are shown in red, LRR repeats in light blue, and the serine/threonine protein kinase domain in green. The amino acids that differ between Gifu and MG20 are highlighted by red boxes. **D**: Sequence alignment of the LRR5 domain in CLE peptide receptors. Amino acids marked in red in *Arabidopsis* PXY/TDR are directly involved in binding the N terminal of the TDIF peptide. A red square highlights amino acid differences between Gifu and MG20. V3 is a conserved region including GxY motif.

### *Pxy* regulates nodule formation

To further investigate the role of PXY during symbiotic interactions, five *LORE1* mutant alleles^64,65^, *pxy-1* to *pxy-5*, were obtained and characterized (**Fig. 5A**). On ¼ B&D medium without nitrate supplement, Gifu plants formed approximately 5 pink nodules 5 weeks post inoculation with *M. loti* R7A, while the *pxy-1* to *pxy-5* mutants developed 2 to 3 pink nodules (**Fig. 5B, C**). The structures of Gifu and *pxy* nodules were similar (**Fig. 5B, C**), but the shoot height in *pxy* mutants was around half that of the Gifu wild type (**Fig. 5D**). In order to further characterize the symbiotic phenotype, we repeated the nodulation experiment using modified Long Ashton medium (**Fig. 5F-H**). The *pxy* nodulation defects were more severe under these conditions, where many white and nearly no pink nodules were formed (**Fig. 5F, G**). To observe the infection status of pink and white nodules, several nodules were sectioned. Both Gifu and *pxy* mutants showed rhizobia colonization of nodule cells (**Fig. 5H**). These results suggest that PXY regulates nodule organ development.

**Figure 5.**
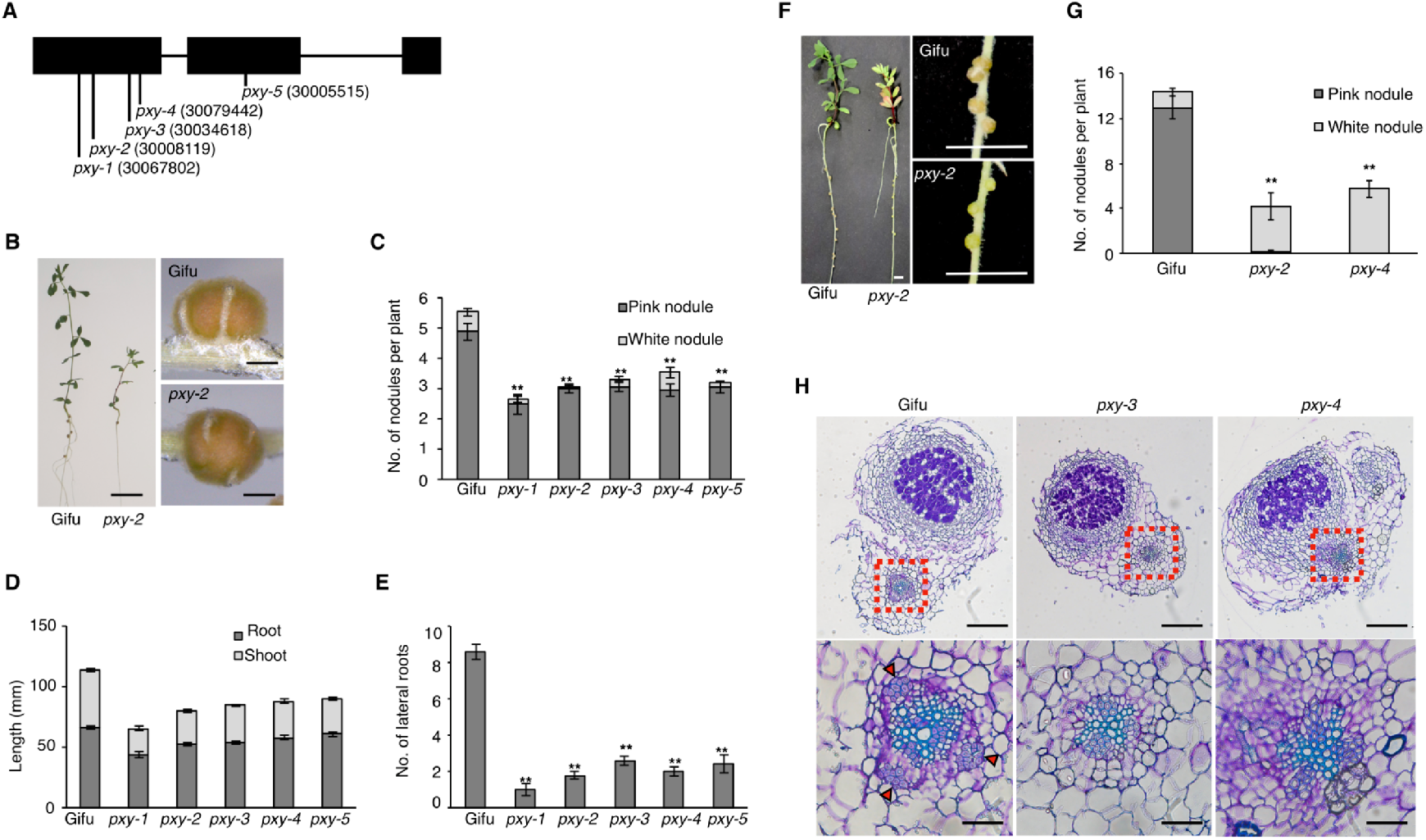
Symbiotic phenotypes of Gifu *pxy* mutants. **A**: Schematic gene model of Gifu *Pxy* showing the positions of *LORE1* retrotransposon insertions. Numbers in parenthesis are *LORE1* mutant line IDs as listed on *Lotus* Base^91^. **B-E**: Symbiotic phenotypes of Gifu wild type and *pxy* plants 5 weeks post-inoculation (wpi) with *M. loti* R7A wild type on 1/4 B&D medium. Bars are 1 cm for whole plants and 5 mm for nodules. **C**: Average number of nodules per plant in Gifu and *pxy* mutants 5 wpi with *M. loti* R7A. **D**: Average length of stem and root in Gifu and *pxy* mutants 5 wpi with *M. loti* R7A. **E**: Average number of lateral roots in Gifu and *pxy* mutants 5 wpi with *M. loti* R7A. **F**-**H**: The Symbiotic phenotype of *pxy* mutants 4 weeks post-inoculation (wpi) with *M. loti* R7A on Long Ashton medium. **F**: Nodule structure of Gifu wild type and *pxy* mutants 4 wpi with *M. loti* R7A. Bars are 0.5 cm. **G**: Average number of nodules per plant in Gifu wild type and *pxy* mutants 4 wpi with *M. loti* R7A. **H**: Cross sections of 2 weeks old white nodules in Gifu or *pxy* mutants inoculated with *M. loti* R7A. Upper panels are whole nodules and bottom panels are closeups of the base of nodules in the red dashed squares. Red arrows point to secondary phloem. Bars are 200 *μ*m in Upper panels, and 50 *μ*m in bottom panels. Two asterisks (p < 0.01) indicate significant differences compared to Gifu plants. *n*= 41 (Gifu), 12 (*pxy-1*), 37 (*pxy-2*), 64 (*pxy-3*), 42 (*pxy-4*), and 24 (*pxy-5*) in **C**-**E**, *n*= 14 (Gifu), 7 (*pxy-2*), and 7 (*pxy-4*) in **G**.

### Vascular differentiation in the *pxy* mutant

PXY functions in cambium proliferation and xylem differentiation^66–68^. In *Arabidopsis, pxy* mutants display a disorganized distribution of phloem and xylem in inflorescence stems and hypocotyls and a decrease in the number of xylem cells^69,70^. In *Arabidopsis pxy* roots, the vasculature appeared normal, but the number of lateral roots was reduced^70,71^ To examine root vascular differentiation in *Lotus pxy* mutants, cross-sections were observed (**Fig. 6**). In the root cross-sections one cm from the root collar, the *pxy* root width was the same as that of Gifu, while the diameter of vascular bundles in *pxy* mutants was significantly smaller than that of Gifu (**Fig. 6A-C**). In order to quantify the abundance of different cell types, the number of xylem and phloem cells were counted (**Fig. 6D**). No significant difference between phloem cell number between Gifu and *pxy* mutants was observed. However, the number of xylem cells was significantly decreased in the *pxy* mutants (**Fig. 6A, B, D**). Interestingly, secondary phloem cells next to the pericycle were properly distributed in Gifu but not in *pxy* mutants (**Fig. 6A, B**). This abnormal distribution of secondary phloem cells in *pxy* mutant was also observed in the root at the base of nodules (**Fig. 5H**). When plants were cultivated in the greenhouse supplemented with nitrate, *pxy* mutants grew normally in the first 6 to 7 weeks, but then displayed growth arrest (**Supplemental figure 2**). In addition, the *pxy* mutants produced fewer lateral roots than Gifu (**Fig. 5E, Supplemental figure 2**). These phenotypic observations indicate that *Lotus* PXY has a function in regulation of root vascular fate and phloem distribution. This is consistent with *Pxy∷GFP* and *Pxy∷triple-YFP-nls* reporter gene expression in vascular bundles in the root meristems (**Fig. 7A, H**). Likewise, *Pxy* expression was detected in cortical cells where nodule and lateral root primordia emerged (**Fig. 7B, C, F, G**), matching the phenotypic defects observed for these organs.

**Figure 6.**
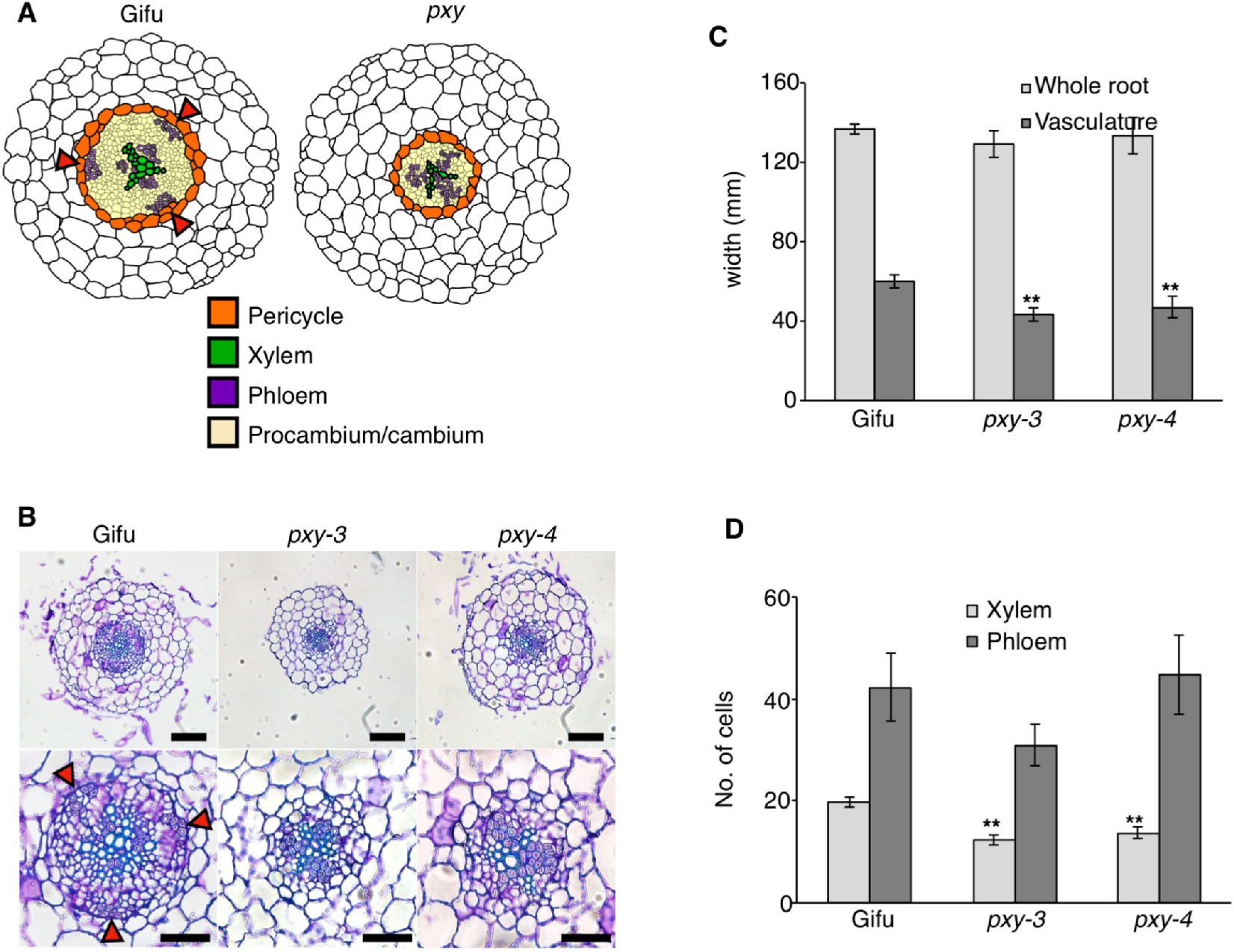
Root vascular organization in *pxy* mutants. **A**, Schematic cross sections of Gifu and *pxy* roots. **B**. Cross sections Gifu, *pxy-3* and *pxy-4* one month old roots. Upper panels show whole root sections, and lower panels show closeups of the vasculature. Red arrows point to secondary phloem. Scale bars are 100 *μ*m in upper panels, 50 *μ*m in lower panels. **C**: Average root widths of Gifu and *pxy* mutants. **D**, Average number of xylem and phloem cells in Gifu or *pxy* mutants. Two asterisks (p < 0.01) indicate significant differences compared to Gifu. *n* = 6 (Gifu), 8 (*pxy-3*), and 8 (*pxy-4*).

**Figure 7.**
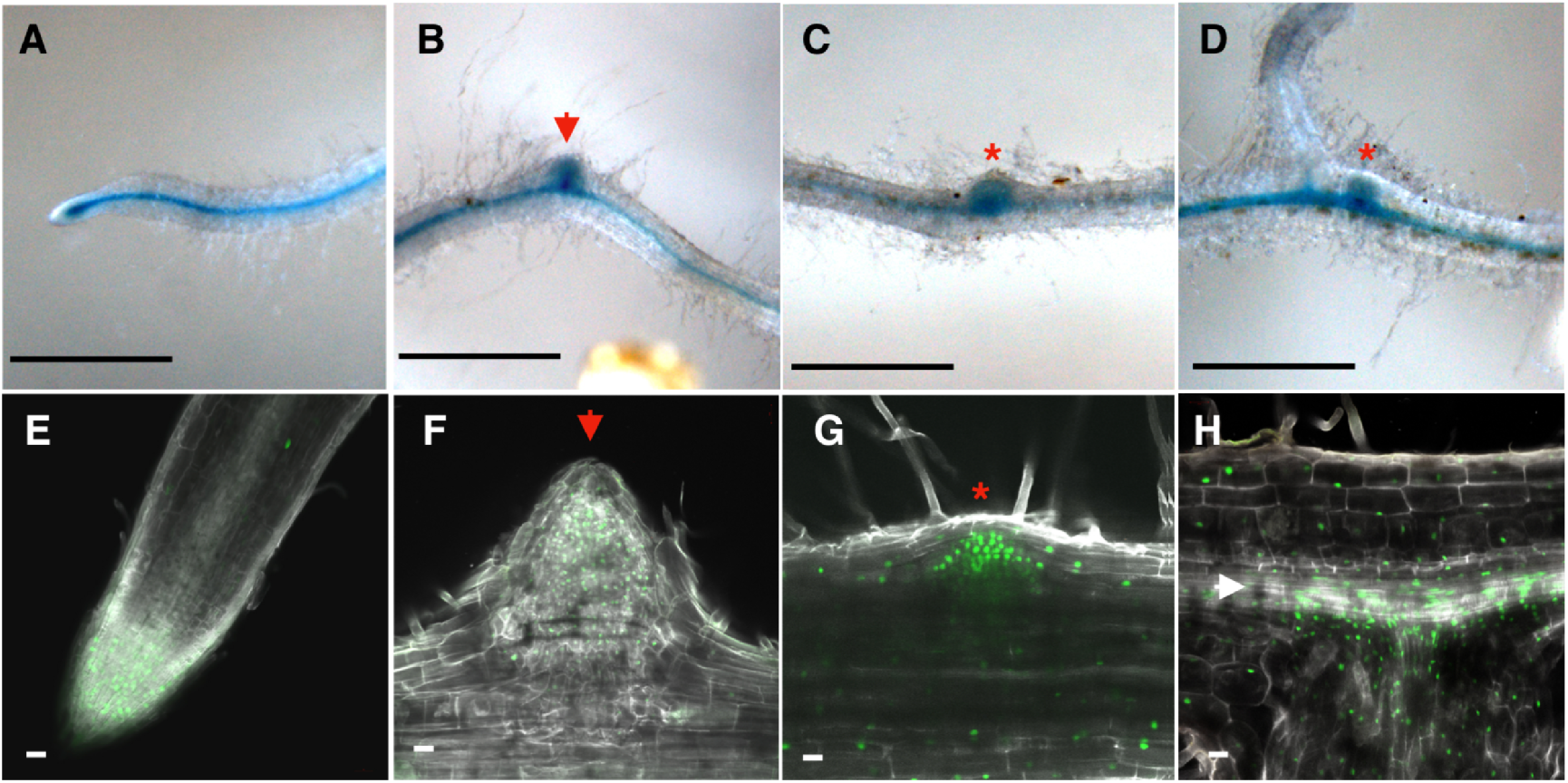
*Pxy* expression in roots. Gifu transformed with a *Pxy∷GUS* (**A** to **D**) or *Pxy∷triple-YFP-nls* constructs (**E** to **H**). **A**, **B** and **E**: Non-inoculated. **C**: 10 days post inoculation (dpi) with *M. loti* R7A. **D** to **H**: 10 dpi with the *exoU* mutant. Red arrows, white arrow and red stars indicate lateral root primordia, vascular bundle and nodule primordia. Scale bars are 2 mm in **A** to **D** and 20 *μ*m in **E** to **H**.

## Discussion

A role for exopolysaccharides in promoting nodulation has been fund in both *Lotus* and *Medicago truncatula*^72^. In *Lotus*, perception of the wild type octasaccharide or the *exoU* truncated pentasaccharide by EPR3 has a positive, respectively negative, effect in Gifu^51,52^. The negative effect of the pentasaccharide is less pronounced in MG20, although there is only a single valine to isoleucine difference in the EPR3 amino acid sequences in Gifu and MG20 (BAI79284.1 and BAI79269.1) (**Supplemental figure 3**). This suggests that the differences between Gifu and MG20 with respect to their *exoU* nodulation phenotype could be due either to differential regulation of EPR3 or to variation in the response downstream of exopolysaccharide perception. In this study, we carried out linkage analysis using F2 and RIL populations to understand the genetic basis of the differential *exoU* response in *Lotus* Gifu and MG20. We identified several genomic regions associated with the phenotype, indicating that the *exoU* response is a complex trait. *Epr3* (chr2.CM0008.630.r2.m, Chr. 2 position 23273436 to 23278205) is not located within any of the candidate intervals, making it unlikely that EPR3 cis-regulatory elements contribute to the intraspecific difference in *exoU* response. Instead, we identified *Pxy* as the gene underlying a major QTL at the end of the chromosome 3. Consistent with the involvement of multiple genes, MG20 *Pxy* did not rescue the Gifu phenotype in transformed roots. However, introduction of MG20 *Pxy* into RI-94 was sufficient to change the symbiotic phenotype and allow pink nodule formation with the *exoU* mutant.

*Pxy* was originally identified in *Arabidopsis* because its loss of function disrupted stem vascular structure, resulting in phloem intercalated with xylem (PXY), while the root vasculature remained unaffected^70,73^. This is in contrast to the *Lotus* root *pxy* phenotype described here, where the vasculature is clearly aberrant in that the number of xylem cells is reduced and the secondary phloem cells are either absent or no longer distinct from the centrally located primary phloem cells (**Fig. 5, 6**). This suggests that the penetrance of PXY function is more pronounced in the complex triarch *Lotus* root than in the simpler diarch *Arabidopsis* root, which has very few files of phloem and xylem cells^74^.

Shortly after its original discovery, PXY was rediscovered as the receptor for the CLE peptide TDIF (tracheary element differentiation inhibitory factor) and acquired the pseudonym TDR (TDIF receptor)^69^. More recently, PXY has emerged as a hormonal signaling hub integrating CLE peptide, ethylene, auxin and cytokinin signaling. PXY suppresses ethylene signaling, and several ERF transcription factors show increased expression in *Arabidopsis pxy* mutants^75^. In auxin and cytokinin signaling, respectively, PXY acts through two similar GSK (glycogen synthase kinase 3) proteins BIN2 (Brassinosteroid-insensitive 2, AT4G18710) and BIL1 (BIN2-like, AT1G06390)^76^. The effects on vasculature phenotype of PXY modulation of cytokinin signaling through BIL1, MONOPTEROS and ARR7/15 were observed in *Arabidopsis* stems^77^. In contrast, reduced formation of lateral roots was the phenotypic basis for the discovery of the PXY effects on auxin signaling through BIN2 and ARF7/19^78^.

Mirroring the *Arabidopsis* phenotype, we observed a prominent decrease in lateral root formation in the *Lotus pxy* mutant (**Fig. 5E, Supplemental figure 2**), indicating a conserved role of PXY in lateral root formation. Interestingly, it has recently been shown that lateral roots and root nodules share a common developmental program and that *ASL18/LBD16a (ASYMMETRIC LEAVES 2-LIKE 18/LATERAL ORGAN BOUNDARIES DOMAIN 16a)* is required for formation of both lateral root and nodule primordia in *Lotus* and *Medicago*^79,80^. In *Arabidopsis*, *ASL18/LBD16a* is under control of Aux/IAA repressors and ARF7/19^81,82^, which are in turn downstream of PXY/TDR. Based on our findings, PXY represents an additional example of a regulator required for both lateral root and root nodule development, which could act upstream of *ASL18/LBD16a*, underlining the molecular genetic overlap between lateral root and root nodule organogenesis pathways.

In *Lotus*, we have shown here that the effect of PXY on nodulation is large enough that natural variation in its coding sequence is sufficient to alter the response to a specific symbiont (**Table 3**). Three SNPs between Gifu and MG20 are located in the PXY coding region (**Fig. 4**) and one of these results in an Arginine (R) to Glycine (G) difference in the super helical structure of the LRR5 motif in the extracellular domain (**Fig. 4D**). The Glycine residue in Gifu is located in a GxY motif that is conserved in other CLE peptide-type receptors in *Arabidopsis* and *Lotus* (**Fig. 4D**). Interestingly, the GxY motif in LRR5 of *Arabidopsis* PXY binds directly to the N-terminal of the TDIF peptide^83–85^. It is conceivable that this Gifu/MG20 polymorphism may result in different TDIF binding affinities for Gifu and MG20 PXY receptors. This, in turn, could influence nodulation signaling and organogenesis, perhaps through effects on ethylene, auxin, and cytokinin signaling, which are all implicated in both nodulation and PXY signaling ^86–88^.

Here, we have identified PXY as a causal component of a nodulation QTL. We have shown that the PXY effect on secondary growth is not limited to stem vasculature, as *Lotus pxy* roots showed clear aberrations in vascular structure. In addition, we found that PXY is required for normal formation of both lateral roots and root nodules. Our work represents a striking example of how natural variation in a nodulation gene can result in a differential symbiotic response, and has provided novel insight into the function of a central regulator of vascular organization, tying it more firmly to root *de novo* organogenesis and highlighting the overlaps between genetic regulation of lateral root and nodule formation.

## Materials and methods

### Plant materials and growth conditions

Plant seeds for analysis of the nodulation phenotype with *M. loti* strains were germinated and grown on ¼ B&D medium or modified Long Ashton medium (**Supplemental table 5**) including 1.4 % agar covered with filter paper in petri dishes. The plant roots were shielded from light by using a metal comb fitting the Petri dish and inserting the lower half of the plates into a rack to keep the root dark. Alternatively, plants were grown in Magenta plastic containers in a 4:1 Leca and Vermiculite mixture supplemented with 60 ml ¼ B&D per container. These plants were inoculated with 500 to 1000 μl *M. loti* culture at OD_600_ = 0.01 to 0.04 per plate and magenta box. Plants were grown at 21 °C under 16/8 h light/dark conditions. Heterologous of *pxy* mutant plants in *Lore1* insertion were grown for harvesting seeds. And homozygous mutants were selected using PCR for each experiment (**Supplemental table 6**).

### Gene and CDS cloning

For the overexpression constructs, the *Xdh2* and *Pxy* genes were amplified from *L. japonicus* Gifu and MG20 genomic DNA by PCR using primers (*Xdh2*; 5’-TCCTA GCTATGAGTTCTCTCA-3’and 5’-CTAGCTAGCATTCCAAGGGA-3’, *Pxy*; 5’-ACCCCAAAACCATGAA CCT-3; and 5’-CATATAAACAGATTAATCAGC-3’) containing attB sites. The *Smc6* CDS was amplified from *L. japonicus* Gifu and MG20 cDNA from root and nodule tissues, which were produced using the 5’-SMART RACE cDNA amplification kit (CLONTECH), by PCR using primers (*Smc6*; 5’-CCGGCGTTTGCAGAATGAAGCGGAGA-3’ and 5’-GAGATTGAACATGATGAAACGCAG-3’) containing attB sites. The PCR amplification products were recombined into the pDONR207 (Invitrogen) using the Gateway BP reaction (Invitrogen) to create entry clone vectors. The entry clone vectors were recombined using Gateway LR reactions (Invitrogen) with the destination vector pIV10∷Ubiquitin_promoter∷GW, to create the constructs: pIV10∷Ubiquitin_promoter∷Gifu_*Xth2*, pIV10∷Ubiquitin_promoter∷MG20_*Xth2*, pIV10∷Ubiquitin_promoter∷Gifu_*Pxy*, pIV10∷Ubiquitin_promoter∷MG20_*Pxy*, pIV10∷Ubiquitin_promoter∷Gifu_*Smc6* and pIV10∷Ubiquitin_promoter∷MG20_*Smc6*. For complement analysis, *Agrobacterium* strain AR1193 was used for hairy root transformation experiments as described previously^62,89^

### Microscopy observation

For expression analysis of a *Pxy∷GUS* and *Pxy∷triple-YFP-nls*, hairy transgenic roots were observed using an Olympus BX53 and a Zeiss Axioplan 2 microscope. For tissue sections, semi-thin sections of nodules and roots were prepared were as previously described^90^. Semi-thin sections were stained by 0.5% toluidine blue and observed using an Olympus BX53 microscope.

### Sequencing library preparation

Tissue was ground in liquid nitrogen to a very fine powder. 2 grams of powder was transferred to 20 ml of ice-cold 1xHB buffer (0.1 M Tris, 0.8 M KCl, 0.1 M EDTA, 10 mM spermidine, 0.5 M sucrose, 0.5%Triton X-100, 0.15 β-mercaptoethanol, pH 9.4-9.5) in a 50 ml Falcon tube. After redissolving by gentle swirling on ice, the suspension was filtered through two layers of Miracloth, by squeezing with a gloved hand. Nuclei were then pelleted by centrifugation with a swinging bucket rotor at 1800 g for 15 min at 4°C. The pellet was resuspended in washing buffer (1xHB buffer without β-mercaptoethanol) and washed 2-3 times. Then the pellet was transferrd into a new 1.5 ml eppondorf tube and resuspended in 500 μl of CTAB buffer (for 200 ml: 4 g CTAB, 16.36 NaCl, 10 ml 0.4 M EDTA pH 8.0, 20 ml 1 M Tris-HCl pH8.0, H_2_O to 200 ml) preheated to 60°C and incubated at 60°C for 30 min with regular mixing. 500 μl chloroform:isoamylalcohol (24:1) was added and contents were mixed by turning the tube until a uniform emulsion was formed. This was followed by spinning at 8000 rpm, 4°C for 10 min and transferring the water phase to a new tube. 5 μl of RNase (10 mg/ml stock) was added and the mixture was incubated at 37°C for 30 min and then put on ice for 5 min. 0.6 volumes of ice-cold isopropanol was added and the sample was, incubated at −20°C overnight. To pellet the DNA, the sample was spun at 6000 rpm, 4°C for 6 min. Finally, the supernatant was discarded an the pellet was washed with 70% ethanol two times. After drying and resuspending in in 55 μl water, the DNA concentration measured using the Nanodrop 1000 instrument. The resulting DNA preparation was fragmented and sequenced by Fasteris (Switzerland) using the standard Illumina protocol for paired-end sequencing. The sequencing reads have been deposited at the Sequence Read Archive (SRA) with accession number (PRJNA623472).

### Analysis of sequencing data

The sequencing reads from both pools were cleaned using TagDust version 1.12 (http://tagdust.sourceforge.net) and reads matching repetitive sequences were removed using custom scripts. The remaining reads were aligned to the MG20 reference genome version 2.5 using bwa version 0.5.9 (bwa aln -t 6 -l 28 -k 2 -I). Variants were called using samtools version 0.1.7 (samtools mpileup with default options). Allele counts were extracted at polymorphic positions and differences in allele frequencies were calculated followed by sliding window analysis using custom perl and R scripts. All scripts are available from https://github.com/stiguandersen/LotusPXY.

### QTL analysis of *exoU* mutant nodulation of Gifu x MG20 RILs

Gifu x MG20 RILs were scored for the average number of nodules formed with rhizobial *exoU* mutants and these observations along with RIL genotypes were used as input for QTL analysis. Data was analyzed using the mqm implementation in the r-qtl package version 1.21-2 (http://www.rqtl.org/). Following data import, data was converted to RIL format (convert2riself), missing genotypes were filled in (mqmaugment), and 1000 permutations were performed to determine significance thresholds (mqmpermutation).

## Author contributions

Symbiotic phenotyping, candidate gene testing, and *pxy* mutant analysis: YK, NS, MK. QTL-seq data analysis: VG, SUA, KS. Project planning and supervision: YK, KS, JS, SUA. Manuscript preparation: YK, SUA, JS.

## Acknowledgements

Thanks to Dr. Yuki Kondo for the discussion of PXY function. This work was supported by the Danish Council for Independent Research | Technology and Production Sciences (10-081677) (SUA), Danish National Research Foundation (DNRF79) (JS), ERC advanced grant (268523) (JS), and Leading Initiative for Excellent Young Researchers (YK).

## Supplemental tables

**Supplemental Table 1.**
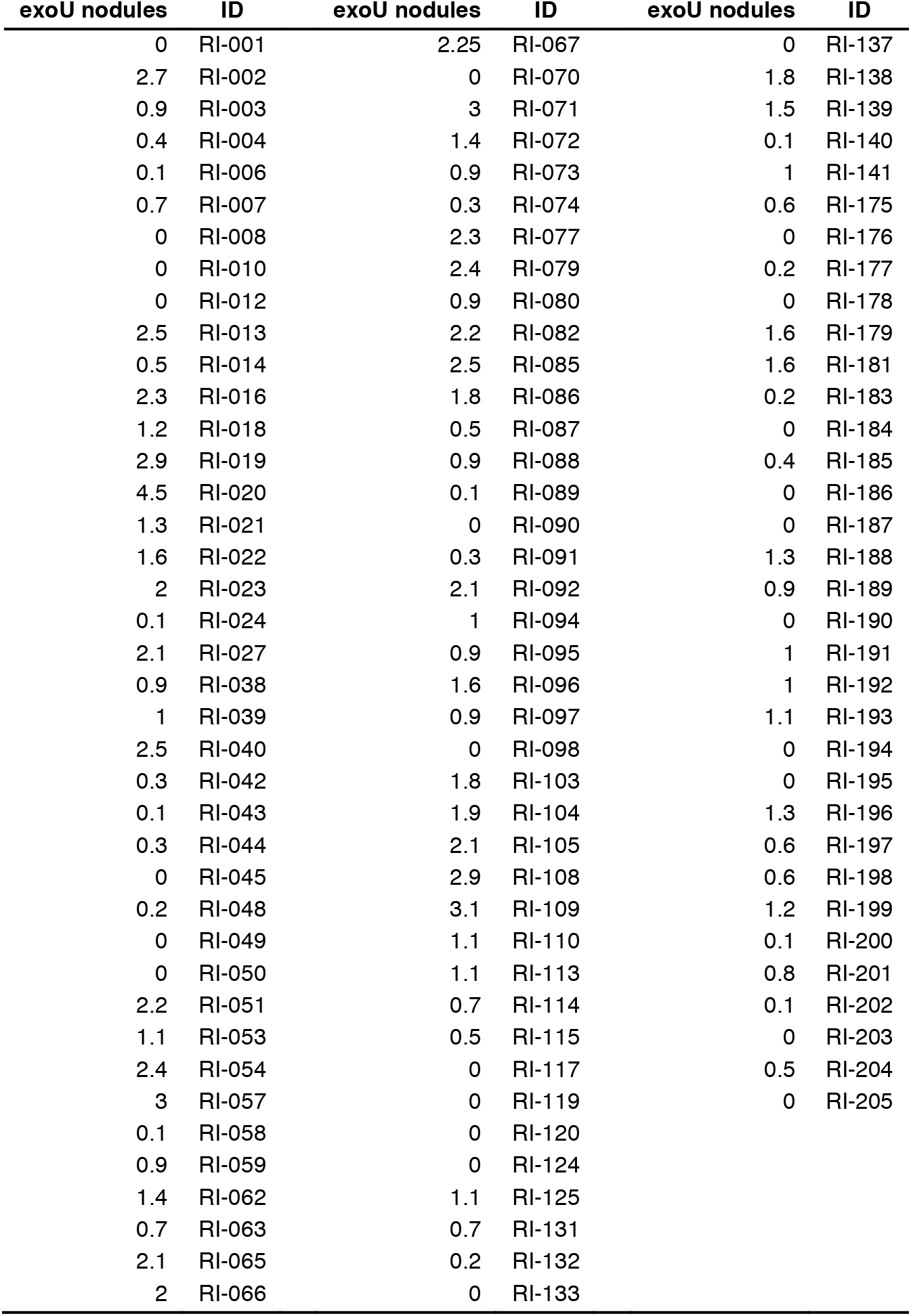
Average number of *exoU* nodules in recombinant inbred lines.

**Supplemental Table 2.**
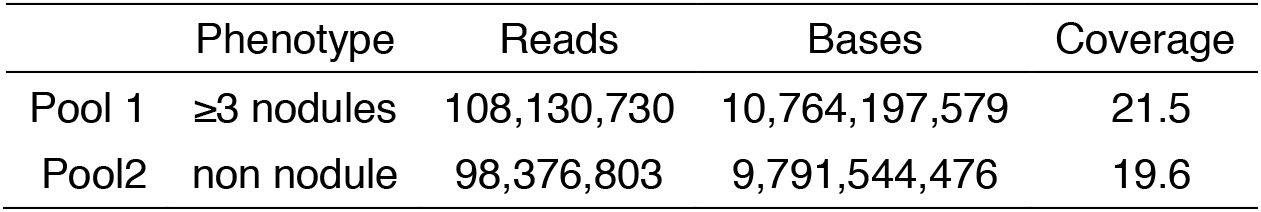
Deep sequencing read depths.

**Supplemental Table 3.**
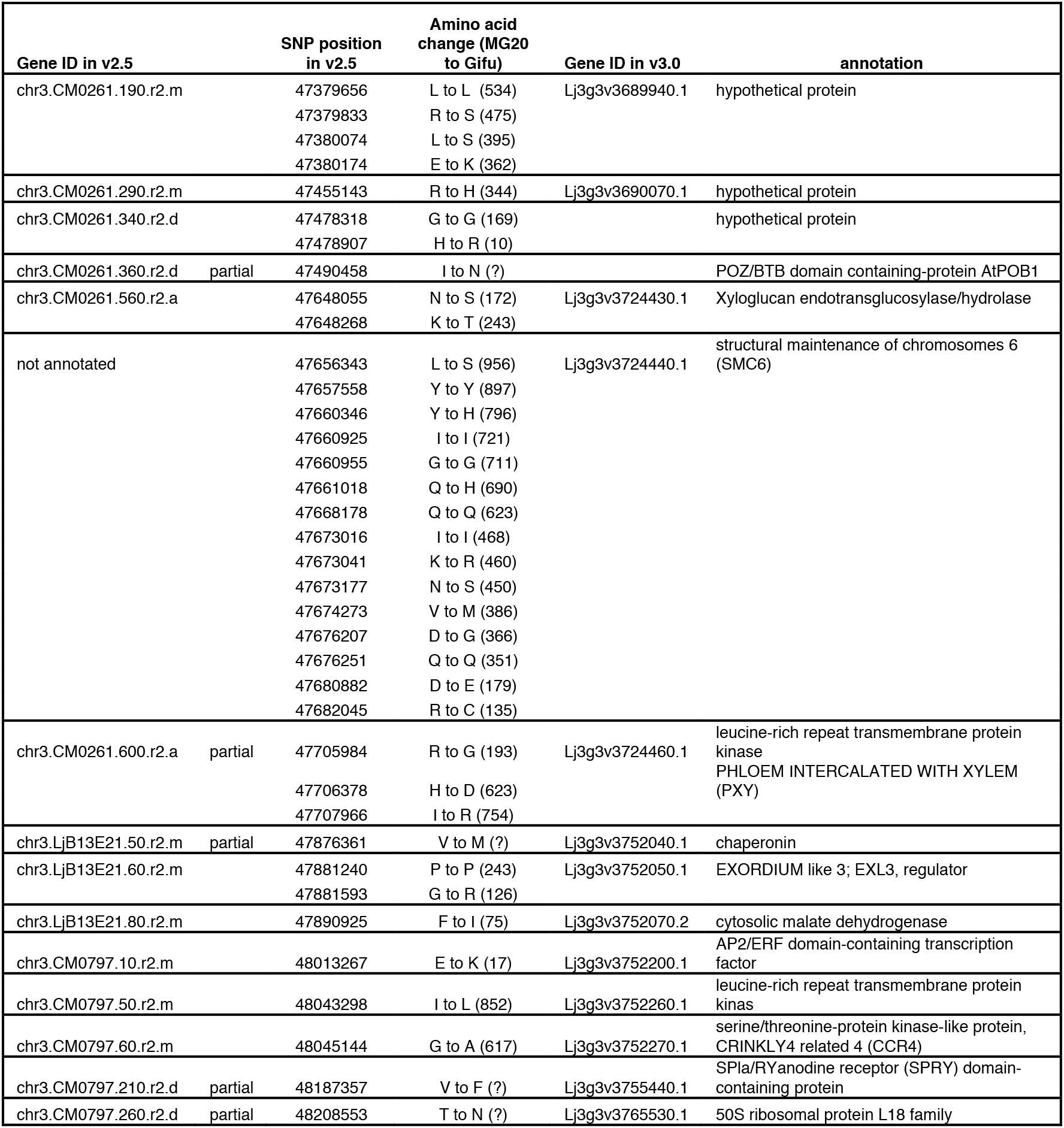
Polymorphisms between Gifu and MG20 for genes located at the end of chromosome 3.

**Supplemental table 4.** Nodule counts for MG20, Gifu, and recombinant inbred line transgenic hairy roots transformed with the genes indicated or an empty vector control (pIV10).

| Plant genotype | Vector | Total no. of plants | No. of plants with nodules | Total no. of nodules | No. of pink nodules | No. of white nodules |
| --- | --- | --- | --- | --- | --- | --- |
| Gifu | pIV10 | 17 | 0 | 0 | 0 | 0 |
|  | MG20 *Xth2* | 29 | 0 | 0 | 0 | 0 |
|  | MG20 *Smc6* | 25 | 2 | 2 | 2 | 0 |
|  | MG20 *Pxy* | 28 | 0 | 0 | 0 | 0 |
| MG20 | pIV10 | 39 | 4 | 4 | 2 | 2 |
|  | MG20 *Xth2* | 27 | 8 | 8 | 8 | 0 |
|  | MG20 *Smc6* | 11 | 2 | 2 | 2 | 0 |
|  | MG20 *Pxy* | 10 | 4 | 5 | 4 | 1 |
| RI-44 | pIV10 | 24 | 0 | 0 | 0 | 0 |
|  | MG20 *Xth2* | 15 | 0 | 0 | 0 | 0 |
|  | MG20 *Smc6* | 29 | 1 | 1 | 1 | 0 |
|  | MG20 *Pxy* | 22 | 0 | 0 | 0 | 0 |
| RI-56 | pIV10 | 10 | 0 | 0 | 0 | 0 |
|  | MG20 *Xth2* | 11 | 0 | 0 | 0 | 0 |
|  | MG20 *Smc6* | 11 | 0 | 0 | 0 | 0 |
|  | MG20 *Pxy* | 6 | 0 | 0 | 0 | 0 |
| RI-91 | pIV10 | 9 | 0 | 0 | 0 | 0 |
|  | MG20 *Xth2* | 6 | 0 | 0 | 0 | 0 |
|  | MG20 *Smc6* | 6 | 0 | 0 | 0 | 0 |
|  | MG20 *Pxy* | 6 | 0 | 0 | 0 | 0 |
| RI-94 | pIV10 | 10 | 0 | 0 | 0 | 0 |
|  | MG20 *Xth2* | 12 | 0 | 0 | 0 | 0 |
|  | MG20 *Smc6* | 18 | 0 | 0 | 0 | 0 |
|  | MG20 *pxy* | 12 | 6 | 7 | 1 | 6 |
| RI-115 | pIV10 | 9 | 0 | 0 | 0 | 0 |
|  | MG20 *Xth2* | 10 | 0 | 0 | 0 | 0 |
|  | MG20 *Smc6* | 20 | 0 | 0 | 0 | 0 |
|  | MG20 *Pxy* | 15 | 0 | 0 | 0 | 0 |
| RI-120 | pIV10 | 13 | 0 | 0 | 0 | 0 |
|  | MG20 *Xth2* | 18 | 0 | 0 | 0 | 0 |
|  | MG20 *Smc6* | 18 | 0 | 0 | 0 | 0 |
|  | MG20 *Pxy* | 12 | 0 | 0 | 0 | 0 |
| RI-140 | pIV10 | 14 | 0 | 0 | 0 | 0 |
|  | MG20 *Xth2* | 14 | 0 | 0 | 0 | 0 |
|  | MG20 *Smc6* | 7 | 0 | 0 | 0 | 0 |
|  | MG20 *Pxy* | 16 | 0 | 0 | 0 | 0 |
| RI-146 | pIV10 | 7 | 0 | 0 | 0 | 0 |
|  | MG20 *Xth2* | 7 | 0 | 0 | 0 | 0 |
|  | MG20 *Smc6* | 9 | 0 | 0 | 0 | 0 |
|  | MG20 *Pxy* | 7 | 0 | 0 | 0 | 0 |

**Supplemental Table 5.** Recipes for ¼ B&D and modified Long Ashton medium.

|  | **B & D** | **Modified**<br>**Long Ashton** |
| --- | --- | --- |
| KNO<sub>3</sub> | | 800 μM |
| KH<sub>2</sub>PO<sub>4</sub> | 500 μM | 750 μM |
| K<sub>2</sub>SO<sub>4</sub> | 1.5 mM |  |
| MgSO<sub>4</sub> 7H<sub>2</sub>O | 250 μM | 750 μM |
| Fe-EDTA | | 100 μM |
| Fe citrate | 10 μM | |
| CaCl<sub>2</sub> H<sub>2</sub>O | 1 mM | 1.1 mM |
| MnCl<sub>2</sub> 4H<sub>2</sub>O | | 10 μM |
| MnSO<sub>4</sub> H<sub>2</sub>O | 1 μM | |
| H<sub>3</sub>BO<sub>3</sub> | 2 μM | 50 μM |
| ZnCl<sub>2</sub> | | 1.75 μM |
| ZnSO<sub>4</sub> 7H<sub>2</sub>O | 0.5 μM | |
| CuCl<sub>2</sub> | | 0.5 μM |
| CuSO<sub>4</sub> 5H<sub>2</sub>O | 0.2 μM | |
| Na<sub>2</sub>MoO<sub>4</sub> | | 0.8 μM |
| KI | | 1 μM |
| CoCl<sub>2</sub> 6H<sub>2</sub>O | | 0.1 μM |
| CoSO<sub>4</sub> 7H<sub>2</sub>O | 0.1 μM | |
| Na<sub>2</sub>MoO<sub>4</sub> 2H<sub>2</sub>O | 0.1 μM | |
| pH | 6.8 | 6.8 |

**Supplemental Table 6.**
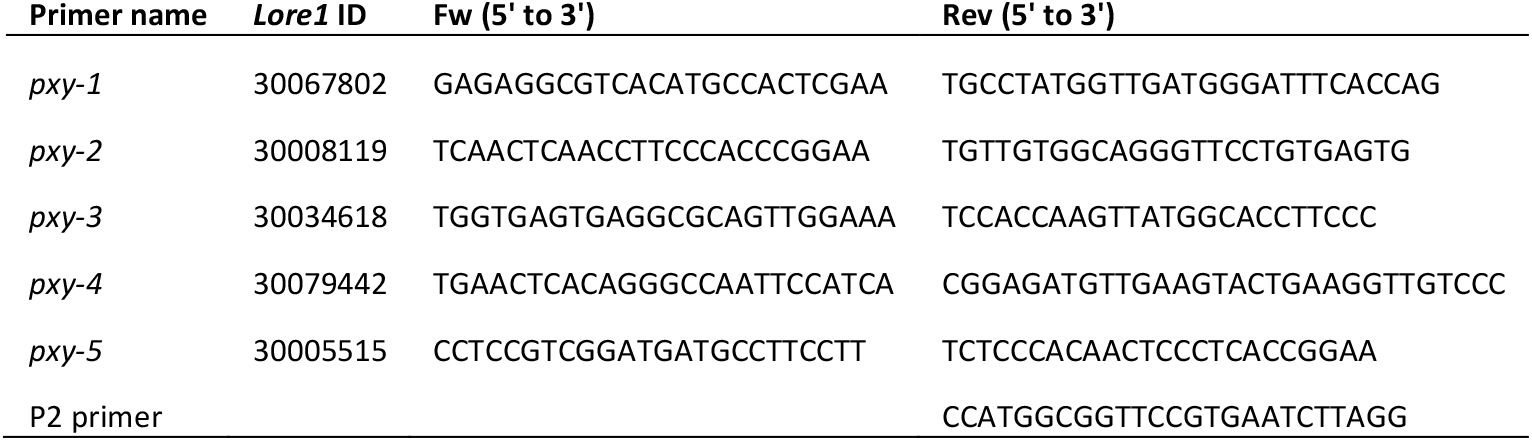
Primers for genotyping of *Pxy LORE1* mutants.

## Supplemental figures

**Supplemental Figure 1.**
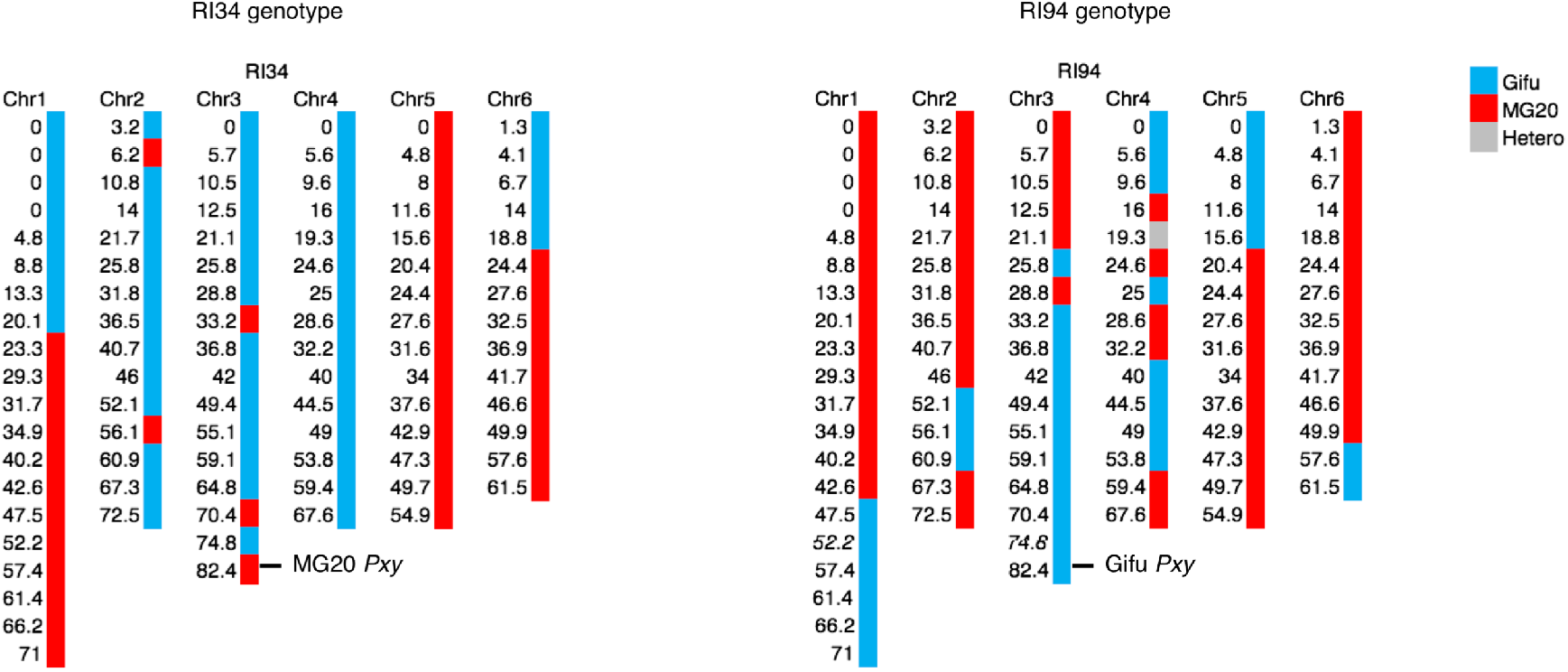
*Lotus* MG20 x Gifu RIL genotypes.

**Supplemental Figure 2.**
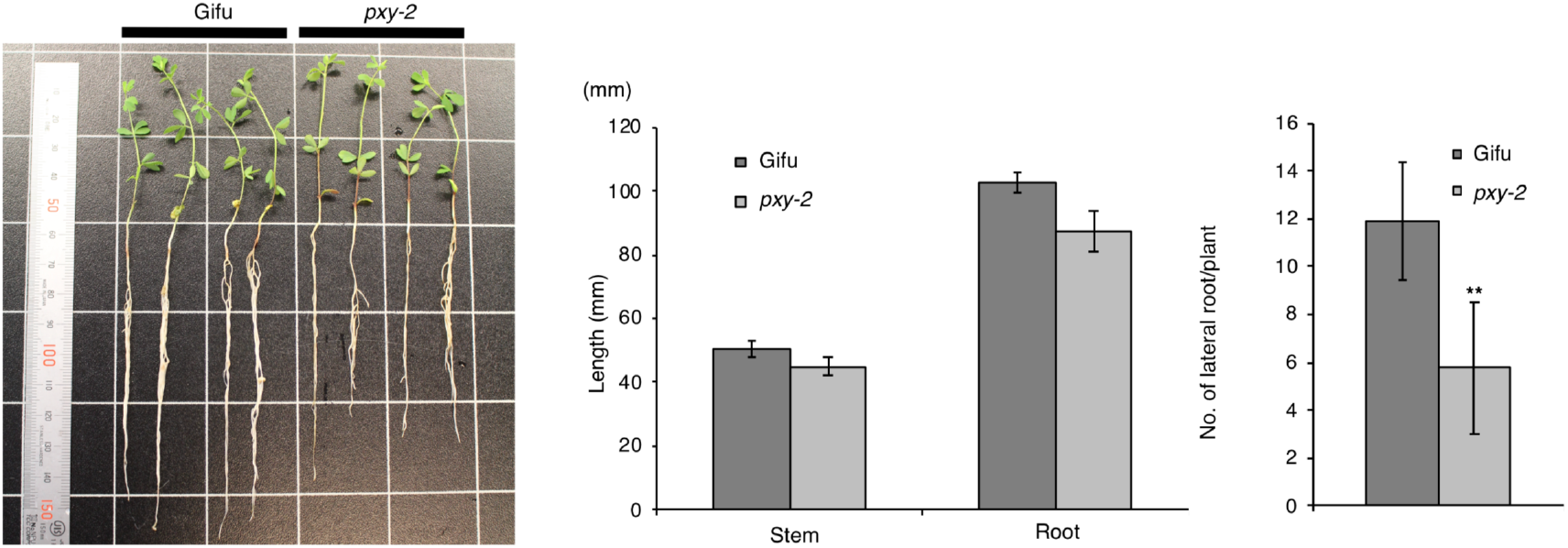
*pxy* mutant phenotypes. Plants were grown with 25 mM KNO_3_ for six weeks after germination.

**Supplemental Figure 3.**
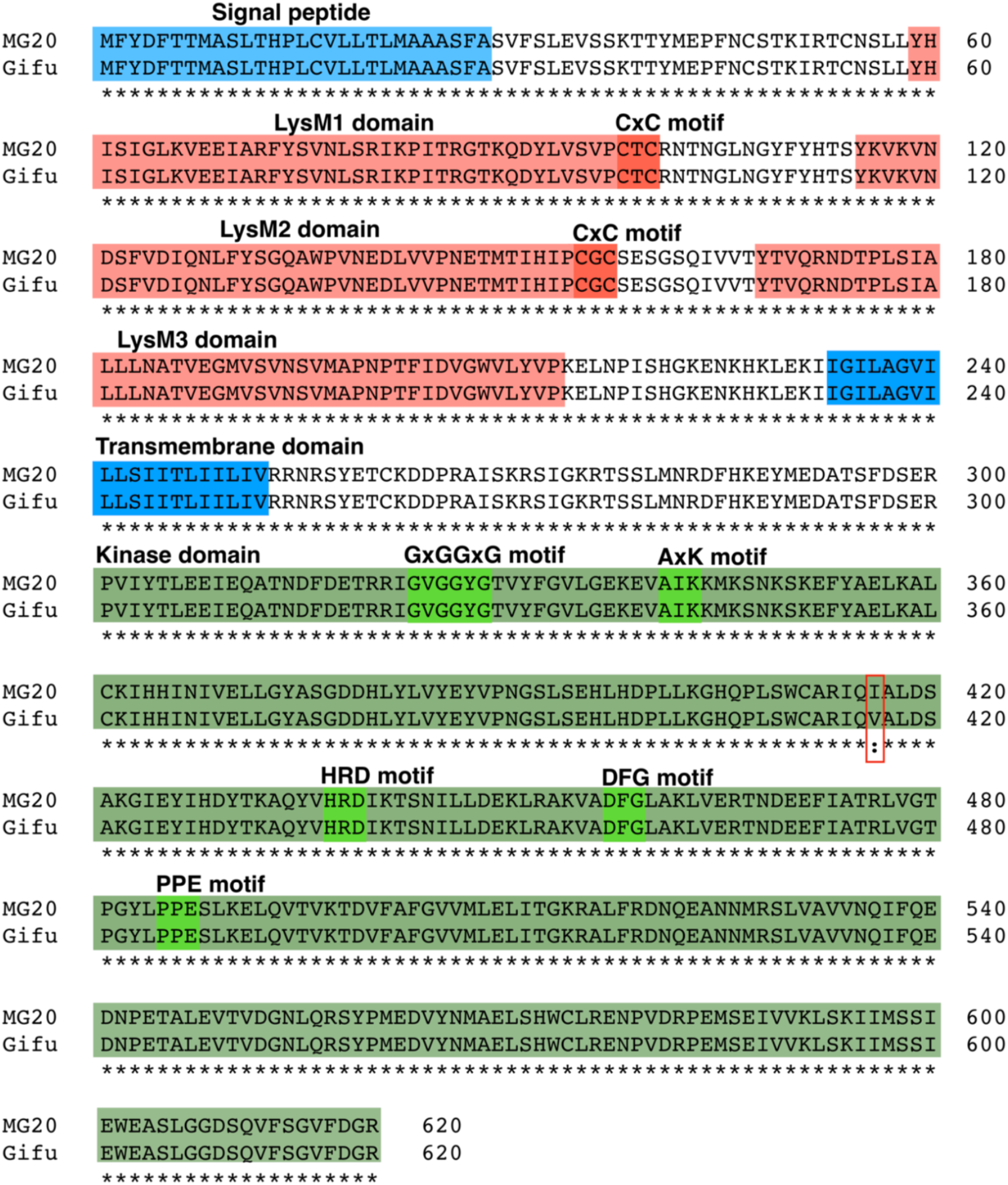
Alignment of EPR3 receptor protein sequences from MG20 and Gifu.

